# Feedback and Feedforward Regulation of Interneuronal Communication

**DOI:** 10.1101/2024.03.22.586312

**Authors:** Oliver Gambrell, Zahra Vahdat, Abhyudai Singh

## Abstract

We formulate a mechanistic model capturing the dynamics of neurotransmitter release in a chemical synapse. The proposed modeling framework captures key aspects such as the random arrival of action potentials (AP) in the presynaptic (input) neuron, probabilistic docking and release of neurotransmitter-filled vesicles, and clearance of the released neurotransmitter from the synaptic cleft. Feedback regulation is implemented by having the released neurotransmitter impact the vesicle docking rate that occurs biologically through “autoreceptors” on the presynaptic membrane. Our analytical results show that these feedbacks can amplify or buffer fluctuations in neurotransmitter levels depending on the relative interplay of neurotransmitter clearance rate with the AP arrival rate and the vesicle replenishment rate, with faster clearance rates leading to noise amplification. We next consider a postsynaptic (output) neuron that fires an AP based on integrating upstream neurotransmitter activity. Investigating the postsynaptic AP firing times, we identify scenarios that lead to band-pass filtering, i.e., the output neuron frequency is maximized at intermediate input neuron frequencies. We extend these results to consider feedforward regulation where in addition to a direct excitatory synapse, the input neuron also impacts the output indirectly via an inhibitory interneuron, and we identify parameter regimes where feedforward neuronal networks result in band-pass filtering.

## I. Introduction

Chemical synapses relay information between two neurons (a presynaptic input neuron and a postsynaptic output neuron) through Action Potential (AP)-triggered release of neurotransmitters. At a fundamental level, this communication is captured via neurotransmitter-filled vesicles that are docked at sites in the active zone of the presynaptic axon terminal, and released when an AP arrives. The depletion of vesicles in response to an AP train is balanced by their replenishment eventually leading to a dynamic equilibrium after a transient depression [1], [2]. Recent work has unmasked the complexity involved in these processes with diverse types of vesicle pools feeding into each other, and differences among vesicles and docking sites in terms of their release and vesicle-recruitment properties, respectively [3]– [12]. This diversity of vesicle pools shapes both short-term and long-term dynamics of neurotransmission in response to high-frequency stimulation of presynaptic neurons [13]–[18].

Characterizing the stochastic dynamics of chemical synapses has been the subject of several works employing both experiments, modeling, and information-theoretic approaches focusing on different sources of randomness and their biological significance [19]–[26]. Our prior work has focused on the stochastic dynamics of docked vesicle counts arising through the coupling of three random processes: random AP arrival, probabilistic vesicle releases, and vesicle replenishment occurring as per an inhomogeneous Poisson process [27]–[29]. This prior work directly coupled the presynaptic vesicle release to changes in postsynaptic membrane potential ignoring the dynamics of the neurotransmitter in the synaptic cleft assuming its instantaneous clearance.

In this study, we explicitly consider the neurotransmitter in the cleft and quantify statistical fluctuations in its levels in terms of physiological parameters. Our results reveal parameter regimes where these fluctuations are minimized at an intermediate frequency of presynaptic AP arrival. A key novelty of this work is to consider feedback regulation (Fig. 1) between the neurotransmitter and presynaptic processes that occurs through sensors known as “autoreceptors” [30]– [32]. Interestingly, our study shows that feedback regulation can play a dual role in both amplifying and buffering fluctuations depending on how quickly the neurotransmitter is cleared from the synaptic cleft relative to the AP arrival and vesicle replenishment rates.

**Fig. 1.**
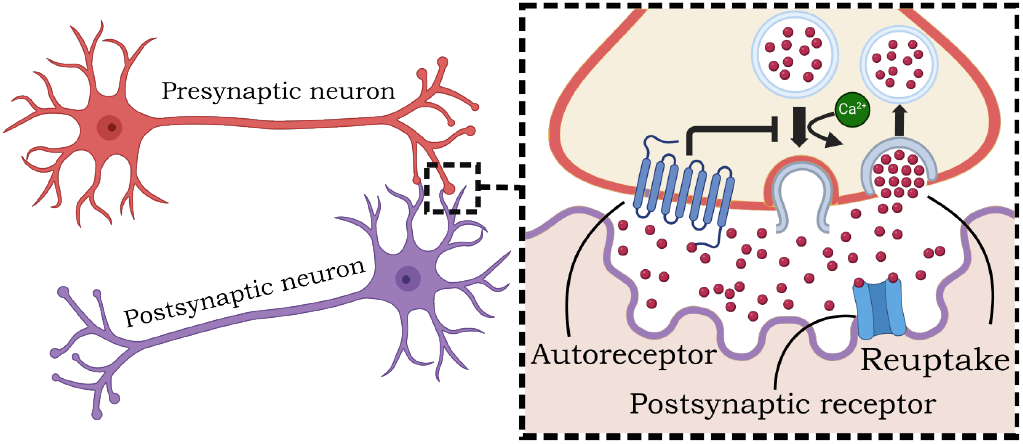
Negative feedback regulation of neurotransmission. Schematic of a presynaptic input neuron forming a chemical synapse with a postsynaptic output neuron. The zoomed-in figure shows docking of vesicles in the presynaptic axon terminal and released neurotransmitters impacting the vesicle docking (replenishment) rate via autoreceptors.

We further connect the neurotransmitter activity to the firing frequency of the postsynaptic neuron and show that while the output frequency typically increases monotonically with respect to the presynaptic input frequency (high-pass filter), in some cases, it can follow a non-monotonic profile leading to a band-pass filter. Finally, we extended this analysis to a network of three neurons that form an incoherent feedforward circuit, and depending on their individual vesicle recruitment capacities the overall system can function as a high-pass or band-pass filter.

## II. Mechanistic modelin of Chemical Synapse

The basic neurobiology of a chemical synapse involves neurotransmitter-filled vesicles present in the active zone of the presynaptic axon terminal. Upon arrival of an action potential (AP) these vesicles are released emptying their content in the synaptic cleft (Fig. 1). These vesicles dock at a fixed number *M* ∈{1, 2, …} of docking sites that essentially defines the upper bound of vesicle counts as determined by the axon terminal’s capacity. Let random processes *n*(*t*) ∈ {0, 1, 2, … *M*} and *x*(*t*) ∈{0, 1, 2, …} denote the number of docked vesicles and the number of neurotransmitters in the cleft at time *t*, respectively. The time evolution of these processes is governed by the following probabilistically-occurring events:

- Upon the arrival of an AP each docked vesicle releases with a *probability of release p*_*r*_ ∈ [0, 1]. Assuming docked vesicles act independently of each other, then conditioned on the number *n*(*t*) of docked vesicles, the number of released vesicles *b* will be a random variable following a binomial distribution

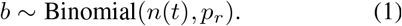
- This evoked vesicle release that reduces *n*(*t*) by *b* leads to an increase in neurotransmitter levels *x*(*t*) by *c*_*x*_ × *b*, where *c*_*x*_ is the number of neurotransmitters per vesicle. The quantities *c*_*x*_ and *c*_*x*_ × *b* are often referred to as quantal size and quantal content, respectively.
- AP-triggered depletion of vesicles is balanced by continuous replenishment of vesicles that occurs with rate *k*(*M* − *n*), where *M* − *n* is the number of empty sites and *k* is the *vesicle replenishment rate* per site.
- Released neurotransmitters are rapidly degraded/cleared from the cleft at a per capita rate *γ*_*x*_, through degradation or uptake by the presynaptic neuron.

In this formulation each docking site is defined by two parameters: the probability of release *p*_*r*_ and vesicle replenishment rate *k*. The site’s proximity to calcium channels can lead to site-to-site differences in *p*_*r*_ and *k*, but here we assume all sites are identical in terms of these parameters. The above events can be rigorously defined through probabilities of occurrences in the next small time interval (*t, t* + *dt*]. For example, assuming APs arrive at the presynaptic axon terminal as per a Poisson process with *input frequency f*, the probability of AP arrival in this interval is *fdt*, and when they arrive, counts are reset as

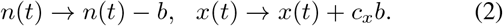

Similarly, the probability of vesicle docking events can be written as

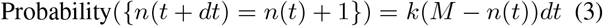

with each event increasing docked vesicle counts by one. Finally, we complete the model description by defining the probability of neurotransmitter clearance

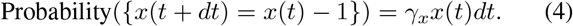

Having defined the presynaptic processes, we define AP firings in the postsynaptic output neuron. The released neurotransmitters impact the postsynaptic neuron’s membrane potential *v*(*t*) as per the linear differential equation

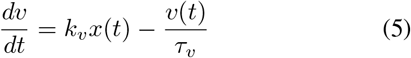

where *τ*_*v*_ is a time constant and *k*_*v*_ is an integration constant quantifying neurotransmitter-mediated opening of receptors for the flow of ionic currents that drive voltage changes. Starting from a resting membrane potential, *v*(*t*) builds up over time to reach a critical threshold resulting in a post-synaptic AP, and *v*(*t*) resets back to its resting potential to repeat the cycle. This model is popularly known as the “leaky integrate-and-fire model” and reduces to a pure integrator for *τ*_*v*_ → ∞ [33]–[36]. Fig. 2 illustrates the time evolution of *n*(*t*), *x*(*t*), *v*(*t*) that function to convert a presynaptic input

**Fig. 2.**
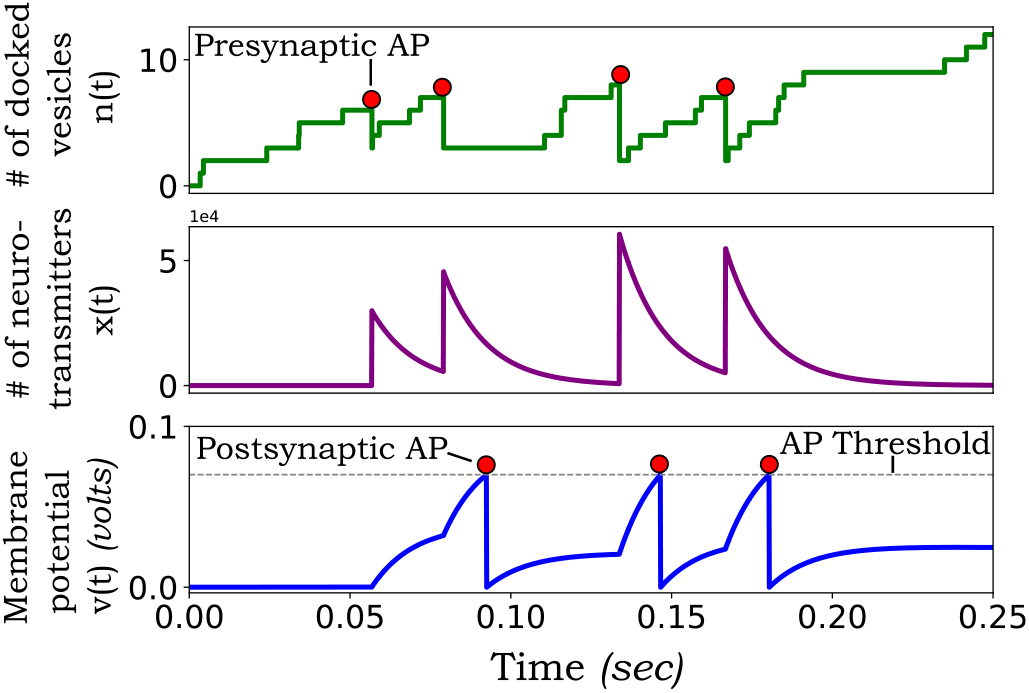
Sample trajectories of the random processes underlying interneuronal dynamics. In response to a train of presynaptic neuron Aps docked vesicles release their neurotransmitter content into the synaptic cleft, which leads to an increase in the membrane potential of the postsynaptic neuron. The top, middle and bottom plots show the sample time evolution of the number of docked vesicles, number of neurotransmitters in the cleft, and postsynaptic membrane potential, respectively. The generation of postsynaptic APs based on the membrane potential reaching a threshold is also shown in the bottom plot. The parameters used are: *M* = 20, *k* = 10*sec*^−1^, *p*_*r*_ = 0.6, *γ*_*x*_ = 100 *sec*^−1^, *k*_*v*_ = 0.0003 *volts, c*_*x*_ = 10000, *τ* _*v*_ = 1 sec, *v*_*th*_ = 0.2 *volts, f* = 20 *H*_*z*_.

AP frequency to a postsynaptic output frequency.

## III. Quantifying statistics of neurotransmission

This section focuses on quantifying the magnitude of statistical fluctuations in *n*(*t*) and *x*(*t*). We start with the model in the previous section and later modify it to consider feedback regulation by making the vesicle replenishment rate a monotonically decreasing function of *x*(*t*).

Before proceeding with the analysis, we discuss some basic nomenclature: the angular brackets ⟨ ⟩ denote expected values, and expectations in the limit *t* → ∞ are represented by 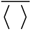. Thus, ⟨*x*⟩ and 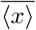denote the average number of docked vesicles at time *t* and at steady-state, respectively. The statistics of count fluctuations are quantified by the steady-state Fano Factors (variance divided by the mean)

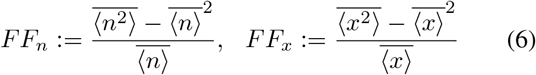

for the stochastic processes *n*(*t*) and *x*(*t*), respectively.

### A. Derivation of moment dynamics

Using the standard tool of moment dynamics for Stochastic Hybrid Systems (SHS) one can derive a system of ordinary differential equations describing the time evolution of statistical moments. Given the probability of occurrences and their corresponding changes in population counts (2)-(4), the time derivative of the uncentered moment ⟨*n*^*i*^*x*^*j*^ ⟩ is

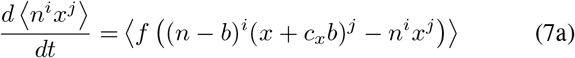

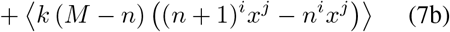

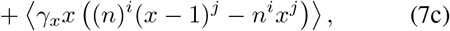

for integers *i, j* ∈ {0, 1, 2, …} [37]–[39]. Intuitively, the right-hand side of (7) is the expected value of the net rate at which an event occurs times the change in the monomial *n*^*i*^*x*^*j*^ when the event occurs. This is then summed across all different classes of events defining the stochastic model.

Taking *i* = 1, *j* = 0 results in the dynamics of the average number of docked vesicles

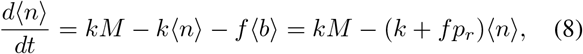

where recall from (5) that *b* is a binomially distributed random variable conditioned on *n* implying

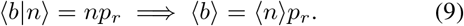

For a given initial condition, (8) quantifies the frequency-dependent synaptic depression in vesicle counts. Similarly, the dynamics of ⟨*n*^2^⟩ is obtained as

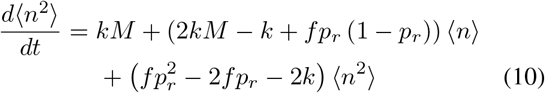

and uses the following properties of a Binomially-distributed *b* to simplify (7) for *i* = 2, *j* = 0

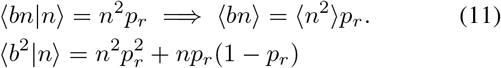

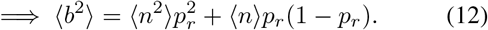

### B. Statistics of the number of docked vesicles

Steady-state analysis of (8) yields

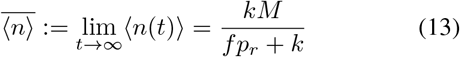

showing a monotonically decreasing average number of docked vesicles with increasing *f* – this is expected given the frequency-dependent deletion of vesicular pools. Similarly, (10) yields the steady-state Fano factor

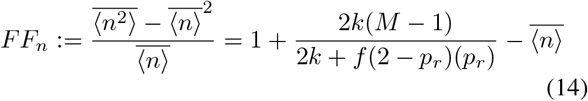

and shows several interesting properties:

- In the limits of low and high frequencies

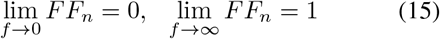

with *FF*_*n*_ = 1 consistent with Poisson statistics.
- If the product *Mp*_*r*_ ≤ 2, then *FF*_*n*_ increases monotonically with *f* to its limit lim_*f*→∞_ *FF*_*n*_ = 1.
- When *Mp*_*r*_ > 2, then *FF*_*n*_ first increases monotonically with *f* to reach a maximum at *f*_*max*_ given by

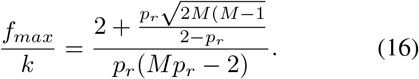

Further increasing *f* leads to the Fano factor decreasing back to 1.
- When *Mp*_*r*_ > 2, the maximum value of *FF*_*n*_ is

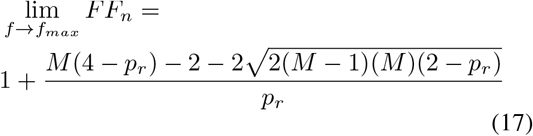

that is approximated by

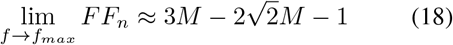

when *M* ≫ 1 and *p*_*r*_ = 1.

The shapes for *FF*_*n*_ as a function of *f* are shown in Fig. 3 with a monotonic (*Mp*_*r*_ ≤ 2) and a non-monotonic (*Mp*_*r*_ > 2) profile. It is important to highlight that Poisson arrival of APs (implying inter-AP arrival times are exponentially distributed with mean 1*/f*) results in super-Poissonian regimes where Fano factors > 1 (Fig. 3). This is in contrast to the case of deterministic arrivals (i.e., time between successive APs are fixed), and in this case Fano factors ≤ 1 are sub-Poissonian across parameter regimes [28], [40].

**Fig. 3.**
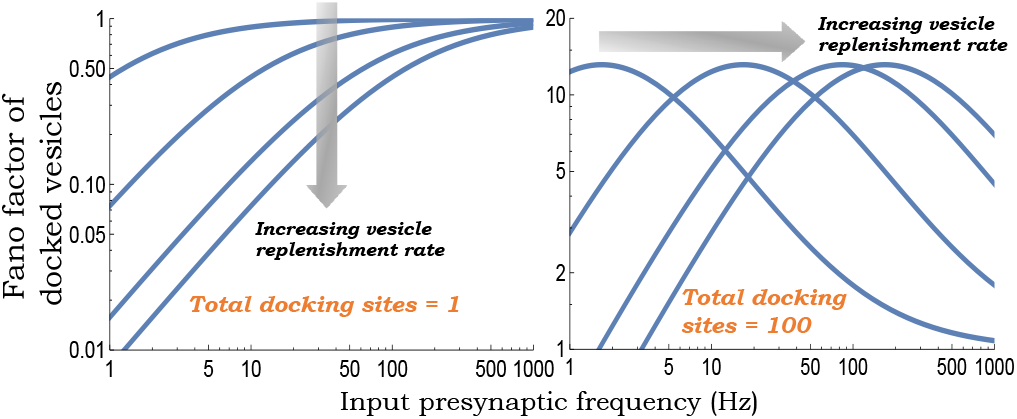
Statistics for the number of docked vesicles can be super or sub-Poissonian depending the number of docking sites. The steady-state Fano factor for the number of docked vesicles as given by (14) is plotted as a function of the AP frequency in the presynaptic neuron. Two different qualitative behaviors are seen: for *Mp*_*r*_ ≤ 2 (left; monotonically increasing with *M* = 1) and *Mp*_*r*_ > 2 (right; *FF*_*m*_ maximized at intermediate frequencies with *M* = 100). The curves are plotted for increasing vesicle replenishment rates *k* = 1, 10, 50, 100 *sec*^*−1*^and the probability of release is taken as *p*_*r*_ = 0.8.

### C. Statistics of the number of neurotransmitters

Taking a similar approach as in the previous subsection, one can derive a system of coupled linear differential equations that describe the time evolution of the moments ⟨*n*⟩, ⟨*x*⟩, ⟨*n*^2^⟩, ⟨*x*^2^⟩, ⟨*xn*⟩. For example, the dynamics of the mean number of neurotransmitters in the synaptic cleft is given by *i* = 0, *j* = 1 in (7) leading to

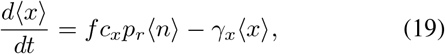

and using (13), yields the steady-state

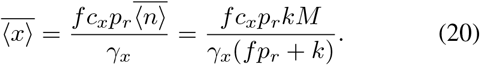

In contrast to (13), 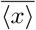 monotonically builds up in the cleft with increasing frequency *f*. Similarly, one can derive an *exact* analytical expression for the Fano factor *FF*_*x*_ that is too complex to be reported here but exhibits the limits:

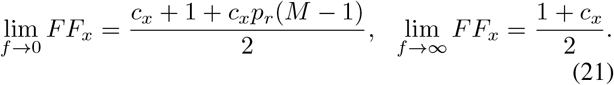

Note *M* = 1 results in the same extreme limits

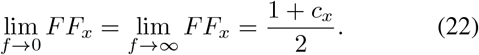

Changes in *FF*_*n*_ with *f* are illustrated in Fig. 4 where depending on *M* the Fano factor profiles are qualitatively different. It is also interesting to contrast these curves with Fig. 3. For example, when *M* = 1, the Fano factor of docked vesicles monotonically increases with *f* (Fig. 3; left), while the Fano factor of neurotransmitter levels is minimized at an intermediate frequency (Fig. 4; left).

**Fig. 4.**
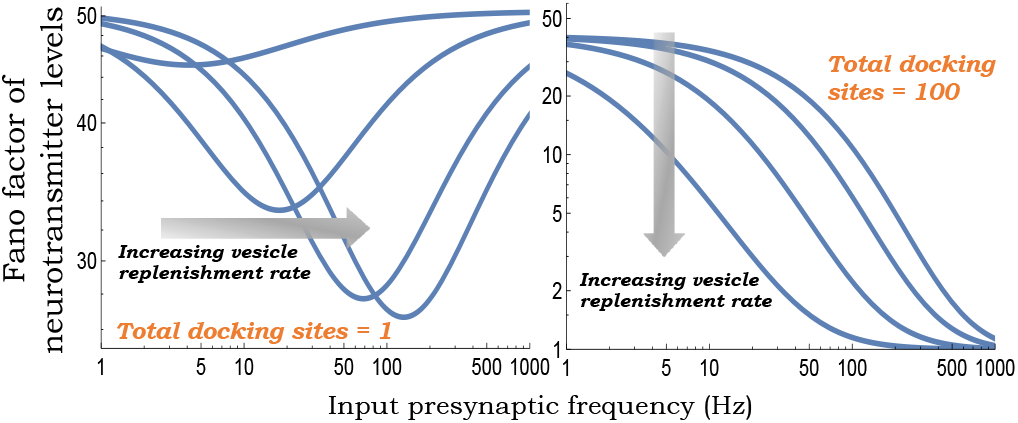
Contrasting profiles of neurotransmitter Fano factor depending on the number of vesicle presynaptic docking sites. The steady-state Fano factor for the number of neurotransmitters in the synaptic cleft as a function of the AP frequency in the presynaptic neuron. For small *M* (left; *M* = 1, *c*_*x*_ = 100), the Fano factor is minimized at an intermediate frequency. For large *M* (right; *M* = 100, *c*_*x*_ = 1), the Fano factor decreases with input frequency in the regime *f* ≤ 1000*H*_*z*_. Other parameters are as in Fig. 3.

### D. Inclusion of feedback regulation

Having systematically quantified the statistical fluctuations in a chemical synapse we turn our attention to the case of feedback regulation. While such feedback can involve the neurotransmitters in the cleft regulating any of the presynaptic parameters *k, M, c*_*x*_, and *p*_*r*_, we specifically consider the case of a regulated vesicle replenishment rate. Towards that end, we modify this rate as *k*(*x*) which is assumed to be a monotonically decreasing function of *x*(*t*).

Given the introduced nonlinearity in the stochastic dynamical system, the classical approach is to consider the Linear Noise Approximation (LNA) [41]–[43] that linearizes the function around the steady state mean

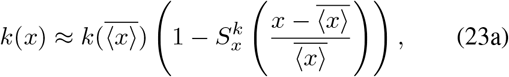

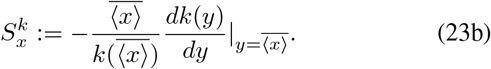

where 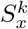 is the log sensitivity of *k*(*x*) with respect to *x* evaluated at its mean. We refer to 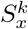 as the feedback strength. The moment dynamics can again be obtained using (7) by replacing *k* (*M* − *n*) with *k*(*x*) (*M* − *n*) that can be approximated as a linear function of *n* and *x* using

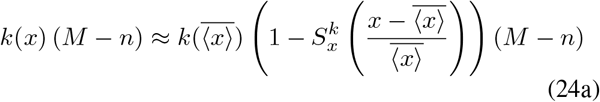

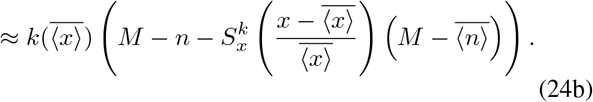

In this small-noise regime, the steady-state mean levels are given as the unique solution to the equation

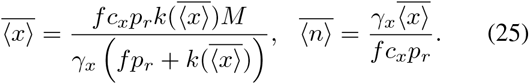

Repeating the moment analysis with the linearly approximated *k*(*x*) results in approximate analytical expressions for all steady-state moments. Fig. 5 (top figure) plots the steady-state Fano factor of neurotransmitter levels with increasing feedback strength as obtained by solving the resulting moment equations at steady-state. Interestingly, negative feedback can amplify or attenuate statistical fluctuation depending on the relative rate of AP arrival as compared to the neurotransmitter clearance rate. This point is further exemplified by plotting the slope (Fig 5; bottom plot)

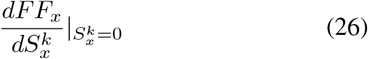

as a function of two dimensionless quantities: 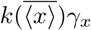 and *fγ*_*x*_ with negative (positive) regions of this plot corresponding to noise attenuation (amplification) with feedback.

**Fig. 5.**
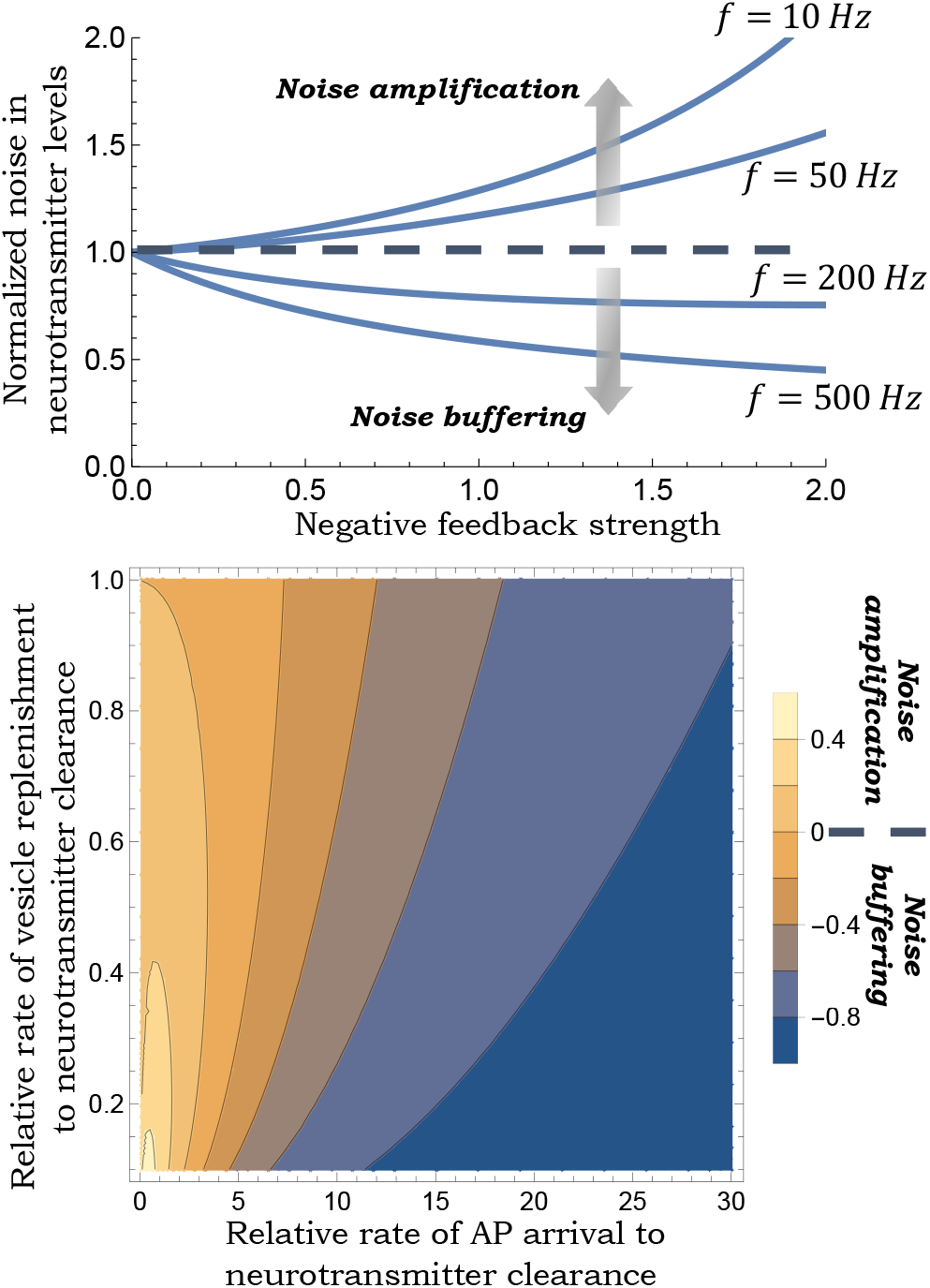
Negative feedback regulation of vesicle replenishment rate can both amplify or buffer statistical fluctuations in released neurotransmitter levels. *Top*: Plot of the steady-state Fano factor *FF*_*x*_ as a function of the negative feedback strength 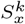. *FF*_*x*_ is normalized by its value at 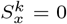. Other <u>para</u>meters taken as *M* = 100, *γ*_*x*_ = 20 *sec*^*−*1^, *c*_*x*_ = 10, *p*_*r*_ = 1 and 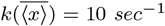. *Bottom*: Plot of the slope (26) in terms of 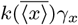 (relative rate of vesicle replenishment compared to neurotransmitter clearance) and *fγ*_*x*_ (relative rate of presynaptic AP arrival compared to neurotransmitter clearance). Other parameters are chosen as in the top figure. Higher clearance rates correspond to the bottom-left region of this plot with positive slope (26) implying that the inclusion of negative feedback increases the magnitude of fluctuations. Similarly, smaller clearance rates correspond to the top-right region of this plot with a negative slope (i.e., feedback inclusion attenuates fluctuations).

## IV. Connecting neurotransmitter activity to postsynaptic Ap firing

We now connect the neurotransmitter levels to AP firings in the postsynaptic neuron. Recall that the generation of these APs is based on the leaky integration of *x*(*t*) as given by (5) that causes an average increase in the postsynaptic membrane potential *v*(*t*). Without loss of generality, we assume the resting membrane potential to be zero. AP firing occurs when the membrane potential reaches a critical threshold voltage *v*_*th*_, and subsequently *v*(*t*) resets back to zero.

Considering the no-feedback case and replacing *x*(*t*) by its steady-state level (20) in (5), voltage integration can be approximated by

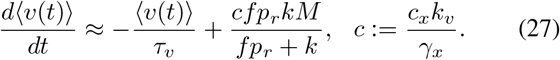

Then, starting from ⟨*v*(0) ⟩ = 0, the average time taken to reach *v*_*th*_ as obtained by solving (27) is given by

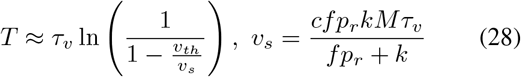

where *v*_*s*_ would have been the mean voltage reached after a long time considering no postsynaptic AP firings (i.e., no resets back to resting potential). Clearly, for (28) to be defined *v*_*s*_ > *v*_*th*_ which leads to result that if

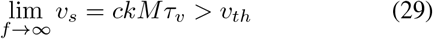

then there exists a minimal frequency *f*_*min*_ such that post-synaptic firing will occur when input frequency exceeds

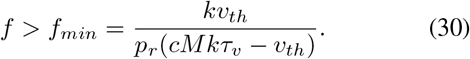

While this analysis is based on a deterministic formulation, in the scholastic formulation even when *f* < *f*_*min*_ (implying that the average membrane potential *v*_*s*_ remains below *v*_*th*_), a noise-induced crossing of the threshold can trigger a post-synaptic AP. However, these noise-induced events become rare as *v*_*s*_ gets much below *v*_*th*_.

Defining the average output frequency

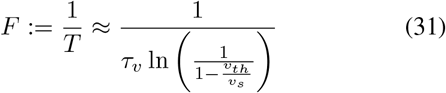

its plot vs. input frequency *f* can be seen in Fig. 6 and reflects high-pass filtering: low input frequencies are filtered out, and *F* monotonically increases with *f* to saturate at

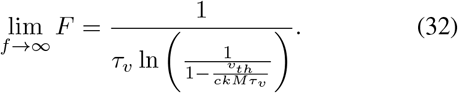

When *ckMτ*_*u*_ ≫ *u*_*th*_

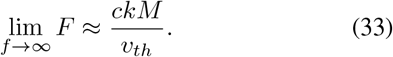

It is important to mention the validity of the approximate formula obtained in (31). Since a postsynaptic AP can only be triggered in response to a presynaptic AP, the average output frequency is always upper bounded *F* ≤ *f* by the input frequency, and (31) is a good approximation only when *F* ≪ *f*. This can be seen, for example in the limit *c* ⟶ ∞, i.e., the number of neurotransmitters per vesicle is so large that a single presynaptic AP triggers a postsynaptic AP. In this case, as *c*→ ∞, *F* → *f*, but the formula (33) would predict an unbounded *F*. This effect can be seen in Fig. 7 which plots the ratio *F/f* as a function of *f* based on the formula (31) and from stochastic simulations. While the formula is accurate for small values of *c* it begins to deviate from its true value for larger values of *c*. Also note from Fig. 7 that this ratio, which can be interpreted as the probability of postsynaptic firing given a presynaptic AP, is maximized at an optimal input frequency.

**Fig. 6.**
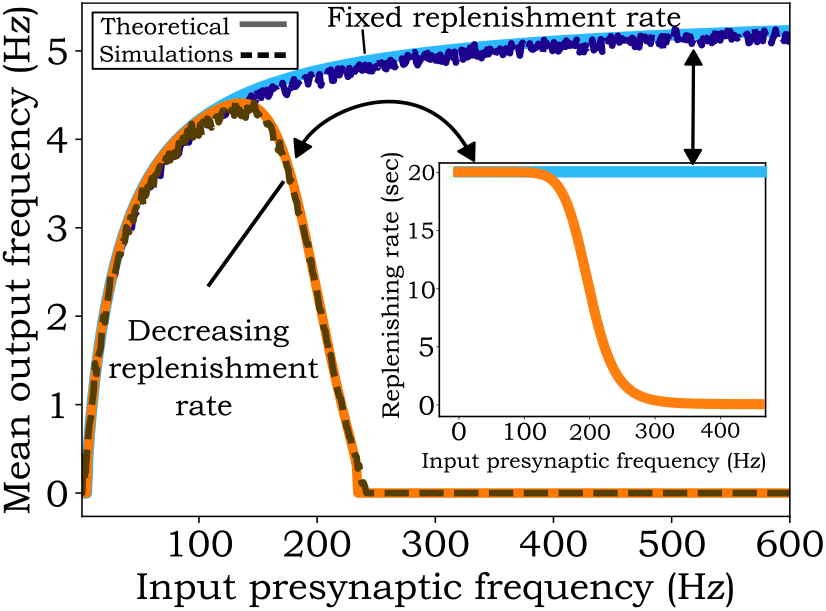
Postsynaptic AP frequency can vary non-monotonically with presynaptic AP frequency depending on the vesicle replenishment rate. The mean output frequency increases monotonically with presynaptic AP frequency *f* for a fixed vesicle replenishment rate (high-pass filter). The plots are shown as per formula (31) that show a good match with their corresponding values from stochastic simulations. When the replenishment rate decreases with increasing *f* (inset figure), then output frequency varies non-monotonically with *f* (band-pass filter). For the band-pass effect, the replenishment rate was take as 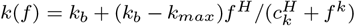 with parameters *H* = 1.7, *k* _*max*_ = 20 sec^*−*1^, *k*_*b*_ = 0 sec^*−*1^, and *c*_*k*_ = 200. The other parameters used in generating this figure are: *M* = 50, *k* = 20 sec^*−*1^, *p*_*r*_ = 0.3, *k*_*v*_ = 0.003 *volts/vesicle, c*_*x*_ = 100, *γ*_*x*_ = 100 sec^*−*1^, *τ*_*v*_ = 5 *sec*, and *v*_*th*_ = 0.2 *volts*.

**Fig. 7.**
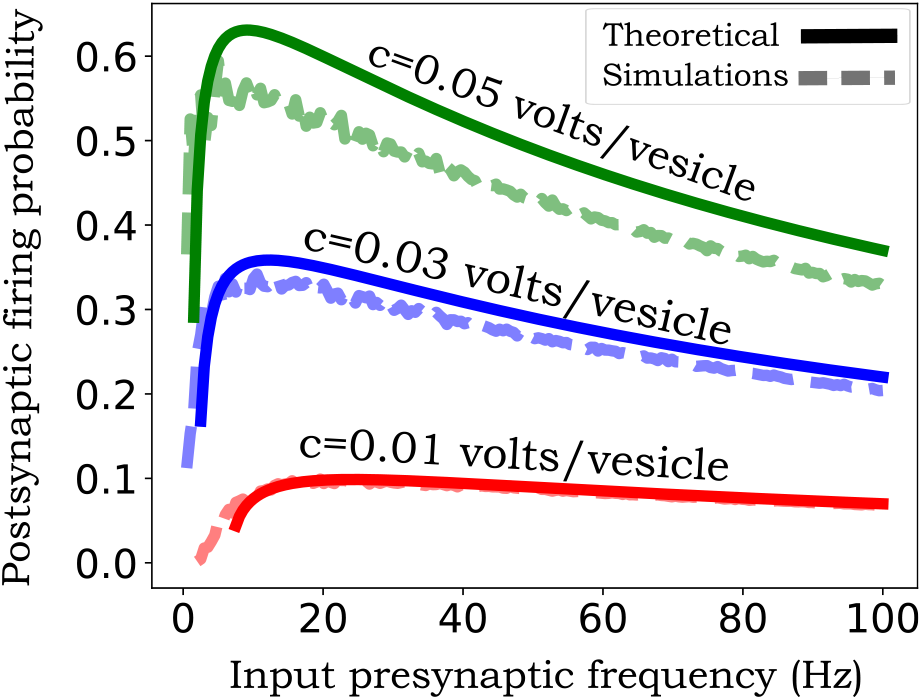
The probability of postsynaptic firing given a presynaptic AP is maximized at an optimal input frequency. The plot of the ratio *F/f* (the probability of postsynaptic firing given a presynaptic AP) as a function of *f* as predicted from (31) and as obtained by running a large number of stochastic simulations. The formula is accurate only in small regimes of the probability of postsynaptic firing. For this plot parameters were taken as *M* = 30, *p*_*r*_ = 0.1, *k* = 10, sec^*−*1^ *c* = 0.02 *volts/vesicle, τ*_*v*_ = 1 *sec, v*_*th*_ = 0.2 *volts, c*_*x*_ = 100, *γ*_*x*_ = 100 sec^*−*1^, and *k*_*v*_ = 0.01, 0.03 and 0.05 *volts*.

In the above analysis, we have assumed all presynaptic parameters (*k, c*_*x*_, *M, p*_*r*_) are constants independent of the input frequency, but in reality they could be frequency dependent. This could occur through frequency-dependent calcium buildup in the axon terminal, or “traffic jams” occurring at high-frequency stimulation impeding neurotransmitter recycling and vesicle maturation processes. Our analysis shows that if the vesicle replenishment rate *k*(*f*) monotonically decreases with *f* then the synapse would function as a bandpass filter – output frequency is maximized at an intermediate input frequency (Fig. 6). Similar results would be seen if the number of neurotransmitters per vesicle *c*_*x*_, or the number of docking sites *M* were to decrease with increasing *f* at high frequencies.

## V. Feed-forward regulation of neurotransmission

We next consider the case of incoherent feed-forward regulation that consists of three components: an excitatory presynaptic (input) neuron, an inhibitory interneuron, and a postsynaptic (output) neuron. The excitatory presynaptic neuron terminates on both the inhibitory interneuron and the output neuron while the inhibitory interneuron terminates on the output neuron (Fig. 8).

**Fig. 8.**
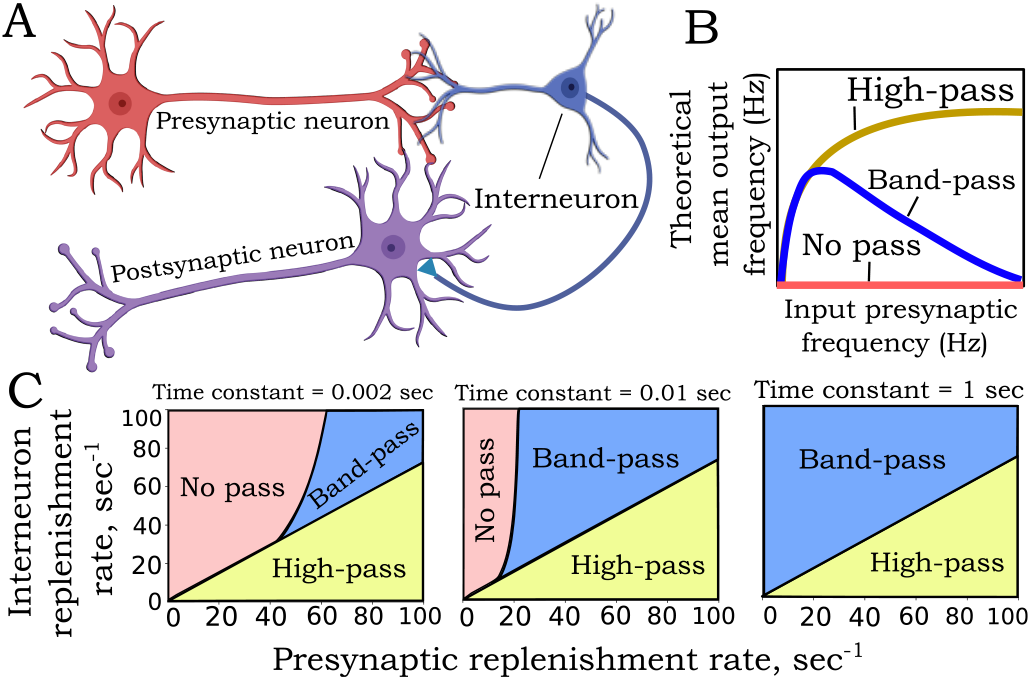
Vesicle replenishment rates of neurons in a feed-forward network can lead to different input-output frequency profiles. **A:** Schematic of an incoherent feed-forward loop, where the presynaptic neuron forms an excitatory synapse with the interneuron and the postsynaptic neuron, while the interneuron forms an inhibitory synapse with the postsynaptic neuron. **B:** Three possible mean output frequency profiles as a function of presynaptic AP frequency *f* : high pass – output frequency increases to a limiting value; band-pass – output frequency is maximized at an intermediate value of *f*, and no-pass – postsynaptic neuron does not to fire. **C:** These regions are shown as a function of the presynaptic and interneuron vesicle replenishment rates for different time constants in the leaky integrate-and-fire model. Parameters used to generate the figure: *c* = 0.01 *volts/vesicle, c*_*i*_ = 0.02 *volts/vesicle, p*_*r*_ = 0.1, *p*_*i*_ = 0.1, *M* = 40, *M*_*i*_ = 50, *v*_*th*_ = 0.2 *volts*.

As we did in the previous section, the mean membrane potential of the output neuron now evolves as

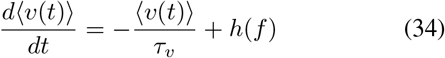

where

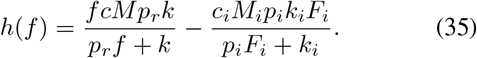

Here *M*_*i*_, *p*_*i*_, and *k*_*i*_ are the number of docking sites, probability of release, and the vesicle replenishment rate, respectively, for the interneuron’s axon terminal forming a chemical synapse with the output neuron. *F*_*i*_ ≤ *f* is the interneuron’s frequency of AP firing, and the parameter *c*_*i*_ is a lumped parameter related to the number of inhibitory neurotransmitters per vesicle, their clearance rate, and their impact on the postsynaptic membrane potential.

It turns out that this overall system can in some parameter regimes exhibit bandpass filtering. We illustrate this point with *F*_*i*_ = *f* (each presynaptic AP will trigger an interneuron AP). These band-pass effects will occur when *h*(*f*) satisfies three conditions:

- *h*(*f*) is an increasing function at low frequencies and the condition

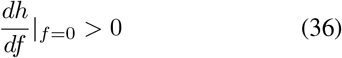

occurs when

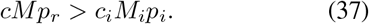
- *h*(*f*) is a decreasing function at high frequencies and the condition

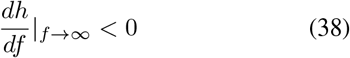

occurs when

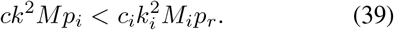
- The maximum value of *h*(*f*) that occurs at an intermediate frequency is higher than *v*_*th*_*/τ*_*v*_ which leads to

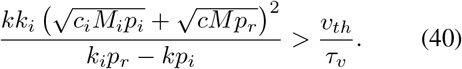

Note that (37) and (39) imply *k*_*i*_*p*_*r*_ > *kp*_*i*_ and (40) is always satisfied for a pure integrator *τ*_*v*_ → ∞.

The region where the incoherent feed-forward circuit works as a band-pass filter, as determined by these inequalities is shown in Fig. 8. In the scenario where *M*_*i*_ = *M* and *p*_*i*_ = *p*_*r*_ these conditions reduce to

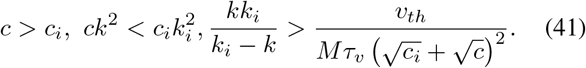

From these inequalities *k*_*i*_ > *k* is a necessary condition for band-pass filtering. Intuitively, this implies that the excitatory neurotransmission gets saturated much faster compared to the inhibitory neurotransmission, and the latter’s inhibitory impact causes the output frequency to decrease with *f* at higher input frequencies.

## VI. Conclusion

Quantifying statistical fluctuations in a chemical synapse is at the core of understanding the fidelity of information flow between neurons. Using a simple yet mechanistic model of a chemical synapse we derived *exact* analytical fluctuations in the levels of the readily-releasable docked vesicle pool and the number of neurotransmitters in the cleft as a function of key physiological parameters.

Our results characterize the changes in the fluctuation statistics as a function of presynaptic input frequency, and in some cases, the Fano factor of released neurotransmitters is minimized at an intermediate frequency (Fig. 4). We considered negative feedback from the released neurotransmitter to the presynaptic parameters with results showing noise attenuation at high input frequencies (Fig. 5). Such high frequencies lead to buildup of neurotransmitters in the cleft, and it is in this regime that the feedback’s role in mitigating fluctuations to prevent further increase in levels may be critical. Our past work investigating autoreceptors in dopamine neurotransmission has also characterized positive feedback loops in neurotransmission that likely kick in at low neurotransmitter levels [32], [44], and would be a part of future work. The usage of LNA for moment computations can also be relaxed in the future by considering moment closure schemes recently developed for Stochastic Hybrid Systems (SHS) [45]–[47], and also using SHS for generalizing inter-AP time intervals to follow arbitrary distributions [48] relaxing the Poisson presynaptic AP arrival assumption.

We connected neurotransmitter levels in the synaptic cleft to the output frequency of the postsynaptic neuron, where the probability of a postsynaptic AP occurring in response to a presynaptic AP can be maximal at an intermediate input frequency (Fig. 7). While we didn’t analyze the noise in postsynaptic AP firings, given that the membrane potential buildup is directly driven by the neurotransmitter levels in the cleft, noise in these AP firings will qualitatively mimic the noise profile of *x*(*t*). Indeed, for a large number of docking sites, our earlier simulations have shown stochasticity in postsynaptic AP firing times to decrease with increasing input frequency [49], [50]. This is consistent with the observation of a decreasing Fano factor in Fig. 4 (right-hand plot). Note here a decreasing Fano factor also implies a decreasing Coefficient of Variation given that the mean neurotransmitter level increases with input frequency.

Finally, we identified two scenarios where neurotransmission can function as a band-pass filter: frequency-dependent reduction in presynaptic parameters, such as the vesicle replenishment rate (Fig. 6) or via an incoherent feedforward loop (Fig. 8). The analysis of the incoherent feedforward circuit was restricted to the interneuron frequency being equal to the presynaptic input frequency, and future work will relax this assumption. Another interesting direction of future work is to use these analytical models of feedforward circuits (both coherent and incoherent) to study several reported invivo examples of such circuits and understand their function in different neurological contexts [51]–[53].

